# Beyond the Forest and the Trees: Overlooking the Overlooked Terrain of Neural State Dynamics

**DOI:** 10.64898/2026.06.04.729738

**Authors:** Tomohisa Asai, Shiho Kashihara, Shinya Chiyohara

## Abstract

State-transition approaches, including EEG microstate analysis and related fMRI methods such as hidden Markov models (HMMs) and co-activation pattern (CAP) analysis, provide widely used tools for coarse-graining neural dynamics into a small set of quasi-stable states. Its utility has been demonstrated across resting-state and task paradigms, with broad applications ranging from cognitive neuroscience to candidate biomarkers for psychiatric and neurological disorders. A fundamental limitation remains, however: nearly all downstream temporal measures are conditional on the template maps defined at the outset. In the conventional pipeline, templates are derived from polarity-invariant clustering of voltage maps at global field power (GFP) peaks, making the resulting state definitions sensitive to preprocessing, sampling, initialization, clustering algorithms, and the choice of cluster number. Consequently, the method captures coarse regularities in EEG dynamics, while only weakly constraining the larger geometric organization from which those states emerge. This template dependence poses a major challenge for reproducibility and for comparisons across studies and EEG caps. Here, we revisit this problem from a topological-geometric perspective. We treat templates not as cluster centroids extracted from GFP-peak maps, but as landmarks embedded in the global structure of a state space constructed from mutual similarities among scalp voltage maps. In this formulation, microstate templates are rediscovered as discrete representatives of dominant axes that organize continuous neural-state topography. This reformulation preserves polarity as a meaningful geometric relation instead of eliminating it at the outset as analytical redundancy. It also shifts attention from isolated state labels to the terrain of the state space itself: the broader relational structure within which local states become interpretable. Using this approach, we show that landmark-based state definitions outperform conventional templates in capturing state structure and improving analytical performance. These findings suggest that the central problem in EEG microstate analysis is broader than clustering optimization: it concerns how to define valid nodes for coarse-graining continuous dynamics without discarding the topology that organizes them. By shifting the conceptual basis of microstate analysis from templates to landmarks, the present approach provides a more principled and potentially more stable foundation for state definition, including in fMRI. This topolo-geometric reappraisal extends conventional microstate analysis and opens a path toward more unified comparisons across datasets, paradigms, and recording systems.

## Introduction

EEG microstate analysis rests on the observation that spatial patterns of scalp potential distributions persist quasi-stably over tens of milliseconds and can therefore be described as a sequence of discrete states (Koenig et al., 1999, 2002; Lehmann, 1971; Lehmann et al., 1987; Michel & Bréchet, 2026; Pascual-Marqui et al., 1995). It can be applied to both resting-state and task-related data, and it provides a concise summary of fast whole-brain dynamics through indices such as the duration, occurrence frequency, coverage, and transition probability of each state. For this reason, it has recently been used not only to advance our understanding of cognitive function (e.g., Michel et al., 2024), but also as a promising source of candidate biomarkers for psychiatric and neurological disorders (e.g., Vass et al., 2025). In addition, the short-term test–retest reliability of microstate measures themselves is generally good (Antonova et al., 2022; Ziogas et al., 2026), and a growing body of evidence supports their high potential for practical application (Asai et al., 2022; Diaz Hernandez et al., 2016; Férat et al., 2025; Tsubaki et al., 2026).

Despite these strengths, the approach has a fundamental unresolved problem. Nearly all derived time-series indices depend strongly on the template states defined at the outset (Michel & Bréchet, 2026). In the current standard procedure, voltage maps at GFP peaks are extracted, clustered using polarity-invariant methods such as modified k-means or AAHC, and the resulting small set of templates is then back-fitted to the entire time series (von Wegner et al., 2018). Although practical, this procedure leaves the representative states vulnerable to sampling, preprocessing, clustering algorithm, initialization, and study-specific implementation choices. Indeed, recent reviews and methodological papers have pointed out that discrepancies in templates make between-group and between-study comparisons difficult to interpret (da Cruz et al., 2020; Michel et al., 2024; Murphy et al., 2023; Trailis et al., 2023), and that approaches such as externally defined meta-microstates or common template schemes have been explored as possible solutions (Koenig et al., 2023).

Template dependence is further complicated by the choice of the number of clusters, k (Custo et al., 2017; Michel & Koenig, 2018; Pascual-Marqui et al., 1995). Although many studies assume four to five canonical microstates, in actual analyses k is often explored using meta-criteria, GEV, cross-validation, and related procedures. Because the optimal k may vary with the data segment, experimental condition, participant group, and noise characteristics, both the number and the shape of templates remain partly indeterminate. Moreover, once the templates are fixed, summary measures such as duration and occurrence frequency are defined on the basis of that particular discretization (Michel & Bréchet, 2026). Conventional microstate analysis therefore offers an effective coarse-grained description of continuous brain dynamics, but the nodes used for coarse-graining remain data-dependent.

Most current approaches are also polarity-invariant. Scalp voltage maps at GFP peaks are typically grouped by polarity-invariant clustering, in which topographies with opposite signs are treated as the same template (Poulsen et al., 2018; von Wegner et al., 2018). This assumption is practically convenient because it treats opposite source orientations as the same state. It may, however, collapse the inversion symmetry of voltage distributions when those distributions are viewed as topographies in a continuous space—that is, as antipodal relations in state space. Conventional methods therefore assign each map to a template before fully exploiting the continuous geometric structure of the full set of voltage maps. The fact that template shapes themselves may differ across groups indicates that the issue concerns state definition itself, rather than implementation alone (Asai et al., 2023).

The present study reframes this problem as one concerning the topographic structure of the state space spanned by scalp maps, rather than the selection of microstate “number” or “templates” (Asai et al., 2022, 2023; Kashihara et al., 2025; Tsubaki et al., 2026). Specifically, we construct a state space from mutual similarities among a broad set of states, not only from conventional GFP-peak maps, and then “rediscover” templates by extracting dominant axes embedded in that space. From this perspective, individual templates become discrete representatives of principal directions in a continuous state space, rather than arbitrary cluster centroids cut from the data. Accordingly, the term “template” is better understood here as a “landmark.” Templates are therefore redefined as quasi-natural nodes derived from the overall topography of the space, rather than as products of a particular clustering initialization (Asai et al., 2023).

This redefinition also allows polarity to be represented naturally. If voltage maps are treated as directional vectors in a state space defined by correlational structure, sign inversion becomes an explicit geometric relation within that space rather than an analytical redundancy. Instead of eliminating polarity in advance, as in conventional approaches (cf., Rošťáková & Rosipal, 2026), it may be more topographically consistent to preserve polarity as an axis of continuous space and coarse-grain the data only when needed. This study is not intended to dismiss EEG microstate analysis. Rather, it repositions the template—from a cluster centroid to an axis of state space—to support a more unified and valid definition of nodes for cross-study comparison (Koenig et al., 2023).

## Materials and Method

In this study, we used large-scale open datasets to embed moment-to-moment brain states in EEG and fMRI (Babayan et al., 2019; Getzmann et al., 2024) into a common low-dimensional state space based on the similarity among spatial patterns, and defined repeatedly occurring locations (stable axes) within that space as topolo-geometric landmarks. Specifically, we constructed a distance matrix from the spatial similarity among neural activity patterns at individual time points and projected it into a three-dimensional space using multidimensional scaling (MDS) (Asai et al., 2023). Within this state space, we defined 14 commonly recurring landmark states (GeoMS14) and compared them with the conventional GFP-peak-based template method (GfpMS14) (see Results for details).

First, we applied different labeling schemes to the same set of EEG/fMRI states and visualized how the resulting states were arranged in the common state space. This allowed us to examine whether GeoMS14 provides a natural partition aligned with the geometry of the state space, whereas template-based methods and existing network labels do not necessarily conform to that structure. We further compared cross-method correspondence and within-method organization using spatial correlations among states, in order to evaluate whether GeoMS14 represents not a mere relabeling, but a reorganization grounded in the geometry of the state space.

Next, we represented the GeoMS14 states as scalp potential distributions in EEG and whole-brain activity patterns in fMRI, and examined whether they were organized as a state series with sign-inversion symmetry and a circular arrangement. On the basis of this structure, we redefined each state as a vector specified by three-dimensional coordinates (x, y, z), and decomposed and reconstructed EEG and fMRI states as linear combinations of orthogonal basis patterns corresponding to each axis. We further extended this basis representation across channel-, voxel-, and ROI-level scales, and examined the correspondence between the state space and functional vocabulary by embedding Neurosynth (Yarkoni et al., 2011) annotation maps into the same space.

Finally, we defined rotational transitions and self-recursive transitions from the state sequences based on GeoMS14 and used them to evaluate individual and group differences. For aging analyses (Kashihara et al., 2025), we compared younger and older groups using the LEMON dataset (Babayan et al., 2019). For sex- and handedness-related analyses, we used the EEG608 dataset (Getzmann et al., 2024; Wascher et al., 2025) to compute rotational direction bias and self-recursive properties. Through these analyses, we evaluated whether GeoMS14 can serve not only as a framework for the static classification of brain states, but also as a basis for describing dynamic indices that capture directionality and cyclicity within the state space.

## Results

First, Figure 1 summarizes the fundamental difference in approach adopted in this study. It is a conceptual comparison between our framework based on topolo-geometric landmarks for defining “common neural states” (A) and the conventional microstate approach based on GFP-peak-based templates (B). The conventional approach defines common states as cluster-center templates; our approach defines them as landmarks in a state space. Templates are intuitive representatives of similar scalp maps, yet they provide limited information about how those maps are positioned within the overall state space or connected through continuous transformations (Shaw et al., 2019). Landmarks are instead defined within a terrain constructed from relationships among all states, allowing continuity itself to remain an object of analysis before the states are assigned discrete labels. Thus, the proposed approach reconceptualizes EEG microstates as shared reference points emerging within continuous neural dynamics, rather than as short-lasting representative maps alone (Mishra et al., 2020).

**Figure 1.**
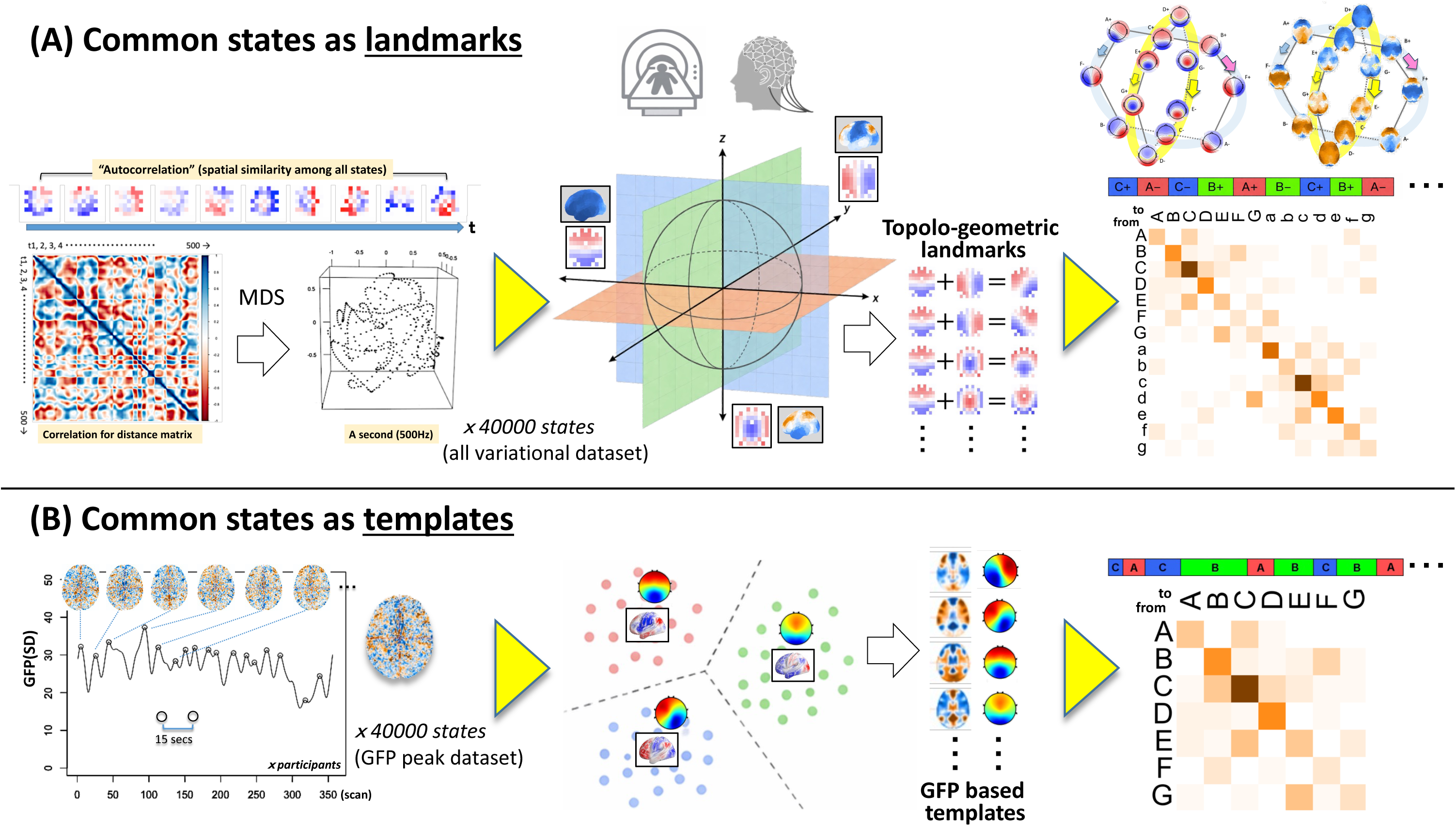
The topolo-geometric reappraisal of neural state dynamics. (A) Newly suggested schematics for depicting neural dynamics both of EEG and fMRI landmarks. (B) Traditional landscaping methodology (non-geometrical) using categorical templates.

In Approach A, the spatial similarity among scalp potential distributions is first evaluated across all states over time, and a state-by-state distance matrix is constructed from these relationships. This distance structure is then embedded into a low-dimensional space, for example using multidimensional scaling (MDS), so that a large number of states can be arranged within a single continuous state space. The key step is to reconstruct the geometric and topological arrangement of the entire state set before partitioning individual states into discrete clusters. As a result, states are distributed within a common spatial terrain, and characteristic positions or directions that repeatedly emerge within that terrain are defined as landmarks shared across participants and time points. These landmarks are common reference points defined as locations in the state space, not just average scalp maps. This makes it possible to treat continuous dynamics naturally, including where individual states arise relative to those landmarks and in what directions they transition. As illustrated on the right side of the figure, each time point can then be assigned to its nearest landmark to obtain a sequence of state labels and a transition matrix, but these labels now carry meaning grounded in the geometry of the entire space. By contrast, in conventional microstate analysis (Approach B), scalp potential distributions at GFP peaks are typically extracted and clustered to define a small number of representative template maps. Each EEG state at a given moment is assigned to the template most similar to its scalp map, yielding a discrete state sequence and transition matrix. In this framework, a common state is a representative map summarizing the distribution within a cluster, and commonality is expressed as an “average center” of similar maps. The template-based approach is therefore well suited to summarizing discrete states, but less suited to representing global structural information, such as relative positions among states and transitions along an underlying continuum. The essential distinction is whether common states are defined as representative patterns (templates) or as shared locations in state space (landmarks). The former tells us what kind of map is typical, whereas the latter tells us what kind of terrain the state set forms and where stable reference points emerge within it. The landmark-based approach can therefore describe common state structure while preserving continuity, directionality, and geometric relationships that are largely lost in conventional discrete classification.

Before comparing the performance of these two approaches, it is important to clarify how they differ at the level of labeling itself. Figure 2 compares how state labels are distributed in the common state space when GeoMS14 (topolo-geometric landmark-based labeling) and GfpMS14 (GFP peak/template-based labeling) are applied to the same set of EEG states. Overall, this figure shows that GeoMS14 organizes EEG states as geometric landmarks within the state space and provides a more consistent spatial positioning of each state, whereas GfpMS14, although effective as a classification into representative templates, is less strongly aligned with the overall terrain of the state space. Put differently, in GeoMS14, labels seem to emerge naturally from the spatial structure itself, whereas in GfpMS14, labels are more likely to behave as categories imposed onto the space afterward. This difference has essential implications for describing the continuous dynamics of EEG states. The three views (View1–3) show the same three-dimensional state space from different angles, with each point representing an individual EEG state. Colors indicate the winner-take-all labels (A± to G±) assigned by each method based on Pearson’s correlation, and representative scalp maps are shown on the right. In the upper row, GeoMS14 produces relatively coherent clusters that occupy stable regions of the spherical state space across views, indicating that its labels correspond to geometrically meaningful positions. In the lower row, GfpMS14 also shows some localization, but the label boundaries and relative positions are more uneven and distorted. Because the point cloud is identical in both rows, the difference comes entirely from how the labels are defined. Thus, although both methods assign the same set of discrete labels, GeoMS14 aligns more naturally with state-space geometry, whereas GfpMS14 behaves more like a post hoc categorization of the same states.

**Figure 2.**
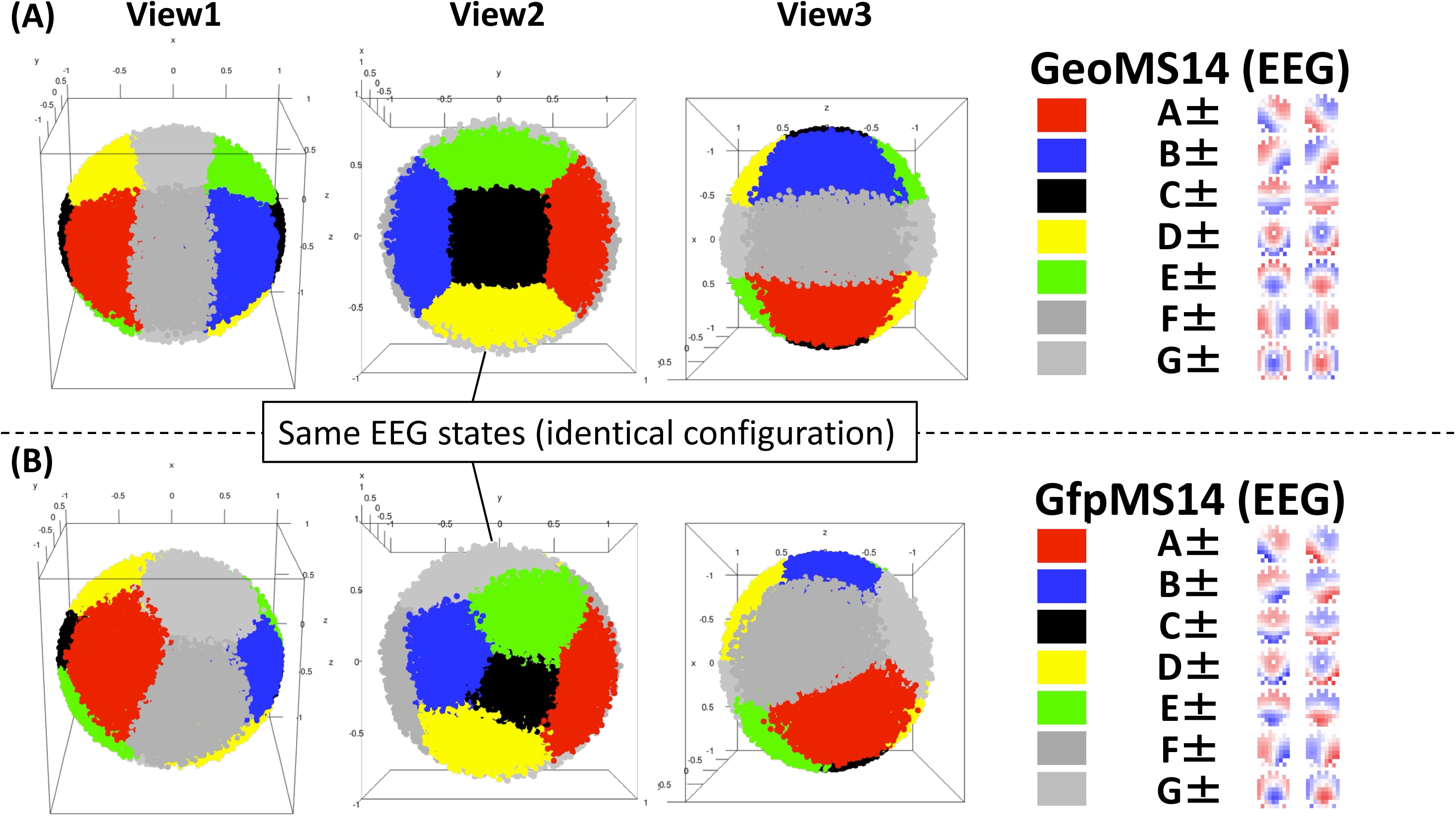
The winner-take-all labels over the instantaneous EEG states on the spherical manifold. (A) Newly suggested 14 templates (geometrical “landmarks”). (B) Traditional GFP-based 14 templates.

Regarding the construction of GfpMS14 and GeoMS14, the GfpMS14 templates were constructed following the logic of conventional GFP-peak-based microstate analysis. First, scalp voltage maps at GFP peaks were extracted and subjected to modified k-means clustering with k = 5. Here, the number of clusters was determined a priori to facilitate comparison with GeoMS14 (see below). This procedure yielded five stable and frequently observed GFP-peak-derived templates, denoted A–E. It should be noted that both the naming and the spatial configuration of microstate classes, particularly those beyond the canonical A–D set, vary across studies and are therefore not directly interchangeable. Two additional templates were then defined by averaging existing templates: F was defined as the average of A and B, whereas G was defined as the average of D and E (for example, F+ was defined as the average of A- and B+). This additional definition was necessary because, in our data, template F/G was not reliably recovered from GFP maxima; this observation is consistent with the broader literature showing that additional noncanonical maps, including vertical topographies, vary across studies and may have ambiguous physiological status (Custo et al., 2017; Jordánek et al., 2025). Rather, it tended to emerge more clearly around GFP minima especially for G (Asai et al., 2023). Finally, polarity was explicitly assigned to each of the seven templates, yielding a total of 14 signed templates, A±–G±. For sign convention, the polarity with a positive frontal field was defined as the positive polarity. However, this criterion did not uniquely determine the polarity of F; therefore, F+ was defined as the polarity in which the left side of the scalp map was positive.

In contrast, GeoMS14 templates were constructed from the geometry of the whole-state space rather than from GFP peaks. A large number of instantaneous EEG states were sampled from the continuous EEG time series without restricting the analysis to GFP peaks; in the present implementation, 40,000 time points were used. Pairwise spatial dissimilarities among these states were computed and embedded into a three-dimensional space using multidimensional scaling (MDS). Because the leading MDS dimensions effectively capture the dominant axes of variance in the state-space geometry, we used the resulting x, y, and z axes to estimate spatial basis patterns. To obtain the spatial basis corresponding to each MDS axis, states were divided into two subsets according to the sign of their coordinate on a given axis. For example, for the x-axis, approximately 20,000 states with positive x-coordinates and 20,000 states with negative x-coordinates were separately collected. Within each subset, individual states were normalized and then averaged. The same procedure was applied to the y- and z-axes. This analysis revealed that the spatial bases corresponding to the three MDS axes were closely related to the signed C, F, and G templates in GfpMS14, respectively. Based on these signed basis templates, the remaining templates were constructed by averaging combinations of C, F, and G. Specifically, A+ was defined as the average of C+ and F−, B+ as the average of C+ and F+, D+ as the average of C+ and G−, and E+ as the average of C+ and G+. The corresponding negative-polarity templates were defined analogously: A− as the average of C− and F+, B− as the average of C− and F−, D− as the average of C− and G+, and E− as the average of C− and G−. The same construction was applied to the fMRI data with arbitrary sign convention (see also Fig. 4 and 6).

Figure 3 compares the spatial similarity among the positive-polarity states (A+–G+) derived from GeoMS14 and GfpMS14 in EEG. Circle size represents the absolute correlation value, and color represents its sign and magnitude, with red indicating positive correlations and blue indicating negative correlations. In (A), spatial correlations are shown for all pairwise combinations of GeoMS14 (rows) and GfpMS14 (columns). Several strong correlations appear in off-diagonal as well as diagonal positions, indicating that the two methods do not show a simple one-to-one correspondence. This suggests that, although both are derived from the same EEG data, they partition the state set differently: GfpMS14 is organized around representative scalp maps, whereas GeoMS14 reflects positions in the state space. In (B), correlations within GfpMS14 show many relatively high positive values, suggesting that several template states remain spatially similar to one another. Thus, separation among states is not always sharp in the template-based solution. In (C), correlations within GeoMS14 appear sparser and more selective. Some states form coherent subgroups, whereas others show strong negative correlations, indicating a more differentiated internal structure. The main point of this figure is that, although both methods define “seven states,” they do not define the same seven states. If they did, panel (A) would show a clear diagonal-dominant pattern. Instead, the presence of strong off-diagonal and negative correlations indicates that GeoMS14 is therefore a distinct state decomposition based on state-space geometry, rather than a simple relabeling of GfpMS14.

**Figure 3.**
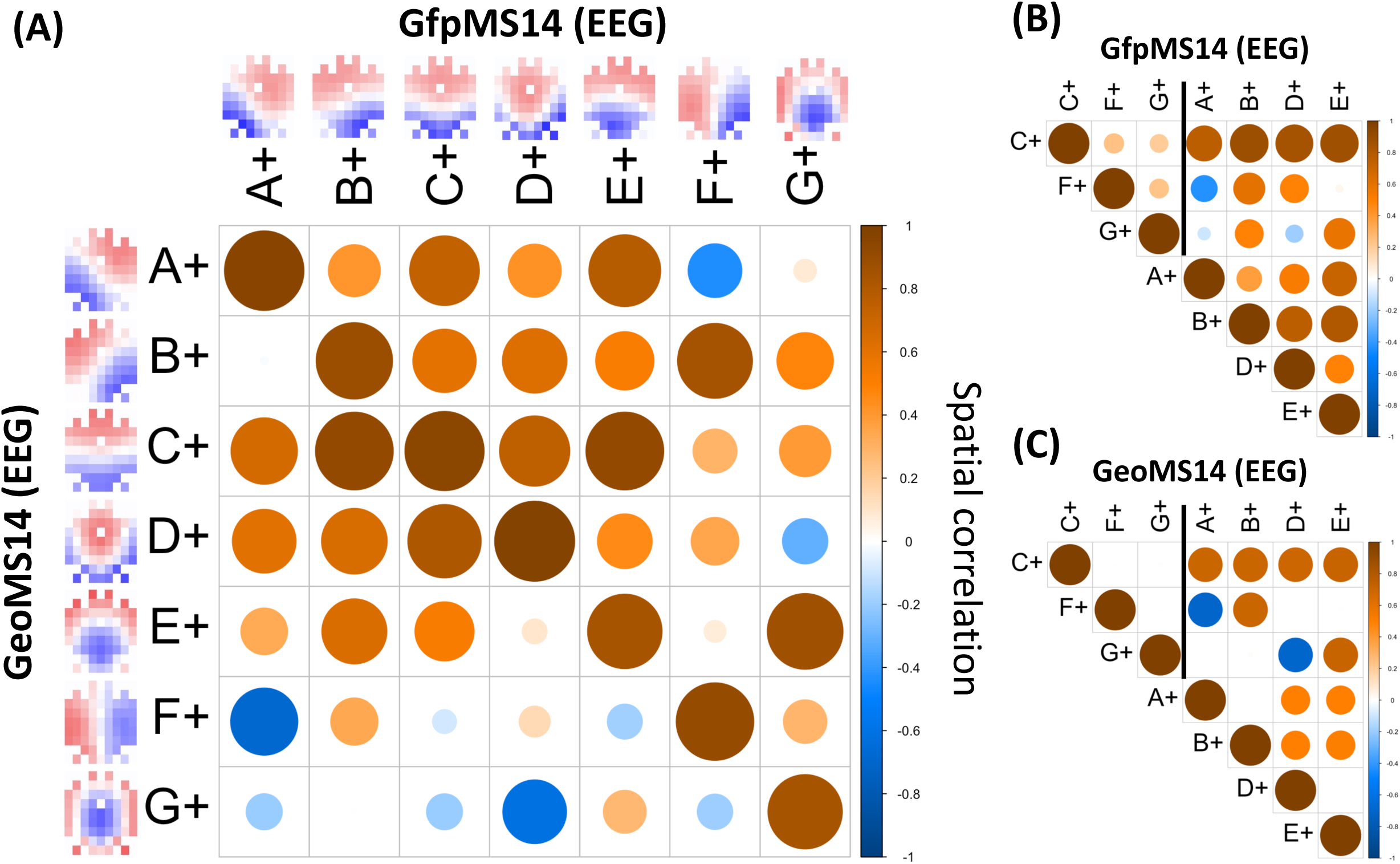
Spatial similarity between/within the new and the old templates in EEG. (A) Between similarity suggests the diagonal high consistency. (B) GFP-based templates correlate with each other. (C) Geometric landmarks keep the fixed metrical distances among them.

The landmark definition can also be applied, in principle, to other recording modalities. Figure 4 compares how the same set of fMRI states is distributed in a common three-dimensional state space under different labeling schemes. In the top row, GeoMS14 forms relatively clear and stable regions in the spherical state space, suggesting that its labels correspond to geometrically meaningful directions or locations. In contrast, the Yeo7 (Yeo et al., 2011) and Yeo14(7±) labels in the middle and bottom rows show much greater overlap and less distinct separation. Although Yeo-based labels are functionally interpretable, they partition this state space less naturally than GeoMS14. Because the underlying fMRI states are identical across all rows, the difference comes entirely from how the labels are defined. Thus, this shows that GeoMS14 is more consistent with the global geometry of the fMRI state space, whereas Yeo labels behave more like externally imposed network categories. This suggests that GeoMS14 provides a more natural framework for describing fMRI states as continuous dynamics. Figure 5 further shows the spatial correspondence between the positive-polarity GeoMS14 states (A+–G+) derived from fMRI and the canonical Yeo7 functional networks.

**Figure 4.**
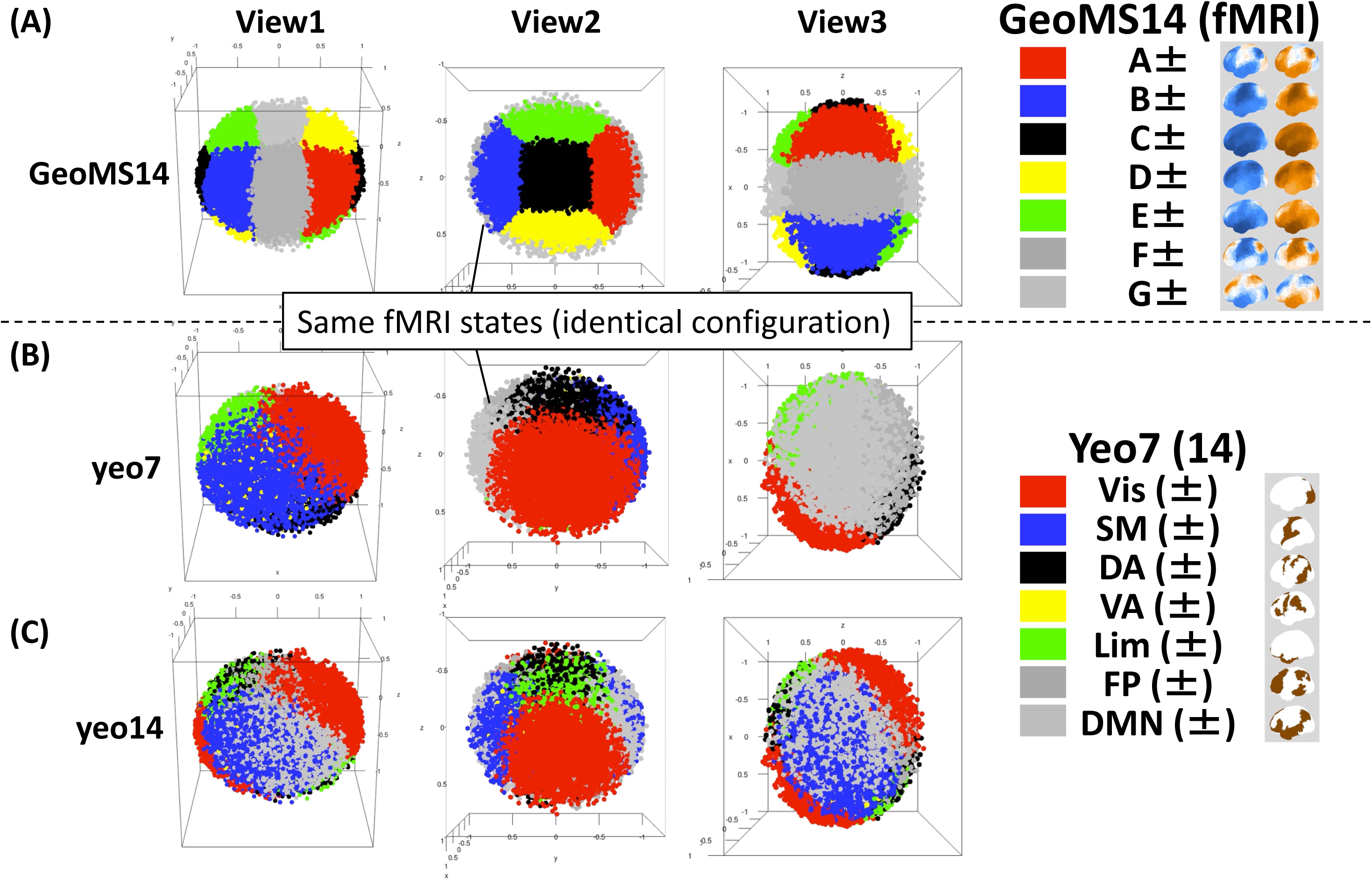
The winner-take-all labels over the instantaneous fMRI states on the spherical manifold. (A) Newly suggested 14 templates (geometrical “landmarks”). (B) Well-known Yeo 7 networks as templates. (C) Those with polarity.

**Figure 5.**
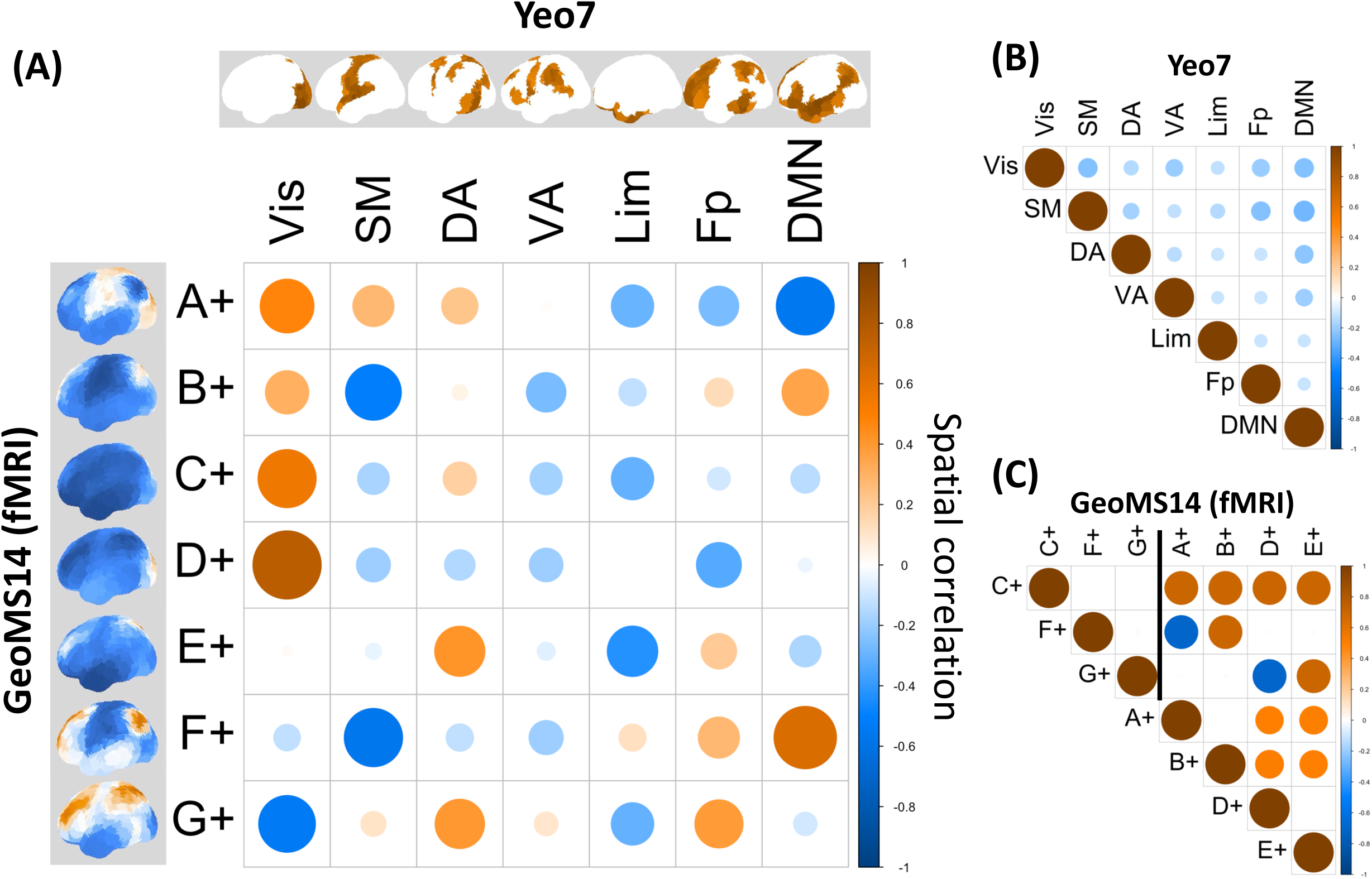
Spatial similarity between/within the new and the old templates in fMRI. (A) Between similarity suggests the functional correspondence of xyz axis (C+ = Vis, F+ = DMN, G+ = CEN). (B) Old templates correlate with each other. (C) Geometric landmarks keep the fixed metrical distances among them like EEG.

In (A), each GeoMS state shows partial positive and negative correlations with multiple Yeo networks rather than a simple one-to-one match. This indicates that GeoMS14 states are not direct equivalents of individual functional networks, but reflect their relative balance and opposition within the state space. In (B), correlations within Yeo7 are generally weak off the diagonal, consistent with a relatively separated network definition. In (C), correlations within GeoMS14 reveal clearer internal structure, including a main positively correlated group and states that are more opposed to others. Overall, Figure 5 indicates that GeoMS14 is related to known functional networks without being reducible to them. Instead, it appears to reorganize those network relationships into a state-space-native landmark representation.

Figure 6 summarizes the overall organization of the GeoMS14 states (A±–G±) in EEG and fMRI. Each node corresponds to one state, shown as a scalp topography in EEG and a whole-brain spatial pattern in fMRI. The arrangement of the nodes reflects both spatial similarity and transition relationships, indicating that GeoMS14 forms a structured state space with intrinsic geometry and preferred transition directions, rather than a mere collection of discrete labels. The small schematics at the top relate the transition probability matrix among states to the classification of rotational transitions. From this matrix, we separately extracted the transitions corresponding to each of the four rotational components. The four rotational transitions and self-recursive transitions together accounted for the majority of all transitions—over 80% in fMRI and over 90% in EEG—although the exact proportions depend on the sampling frequency. Thus, the large circular arrangement reflects a higher-order organization that integrates spatial correspondence among states with their directional transition structure. Across both EEG and fMRI, the states are arranged from A+ to G+ and from A− to G− in a polarity-symmetric and approximately circular configuration (sign convention is arbitrary, therefore could be opposite), making both sign-inversion symmetry and cyclic relationships visible. The same global organization appears in both modalities. Although the EEG and fMRI patterns themselves are different, their states follow a similar ordered structure, suggesting that GeoMS14 reflects a more general topological principle of neural state organization rather than a modality-specific artifact. The central message of this figure is that GeoMS14 is best understood as a landmark system defined on a structured state manifold, rather than as 14 isolated states. Its significance lies in identifying recurring states and showing that they are organized by continuity, symmetry, and directional transition structure. Figure 7 visualizes the GeoMS14 states in fMRI as cortical spatial patterns and shows their relationships from multiple viewpoints. The four panels correspond to the left lateral, right medial, superior, and posterior views, respectively. Each node represents one GeoMS state; gray lines indicate dominant rotational trajectories (horizontal and vertical, see below) among states.

**Figure 6.**
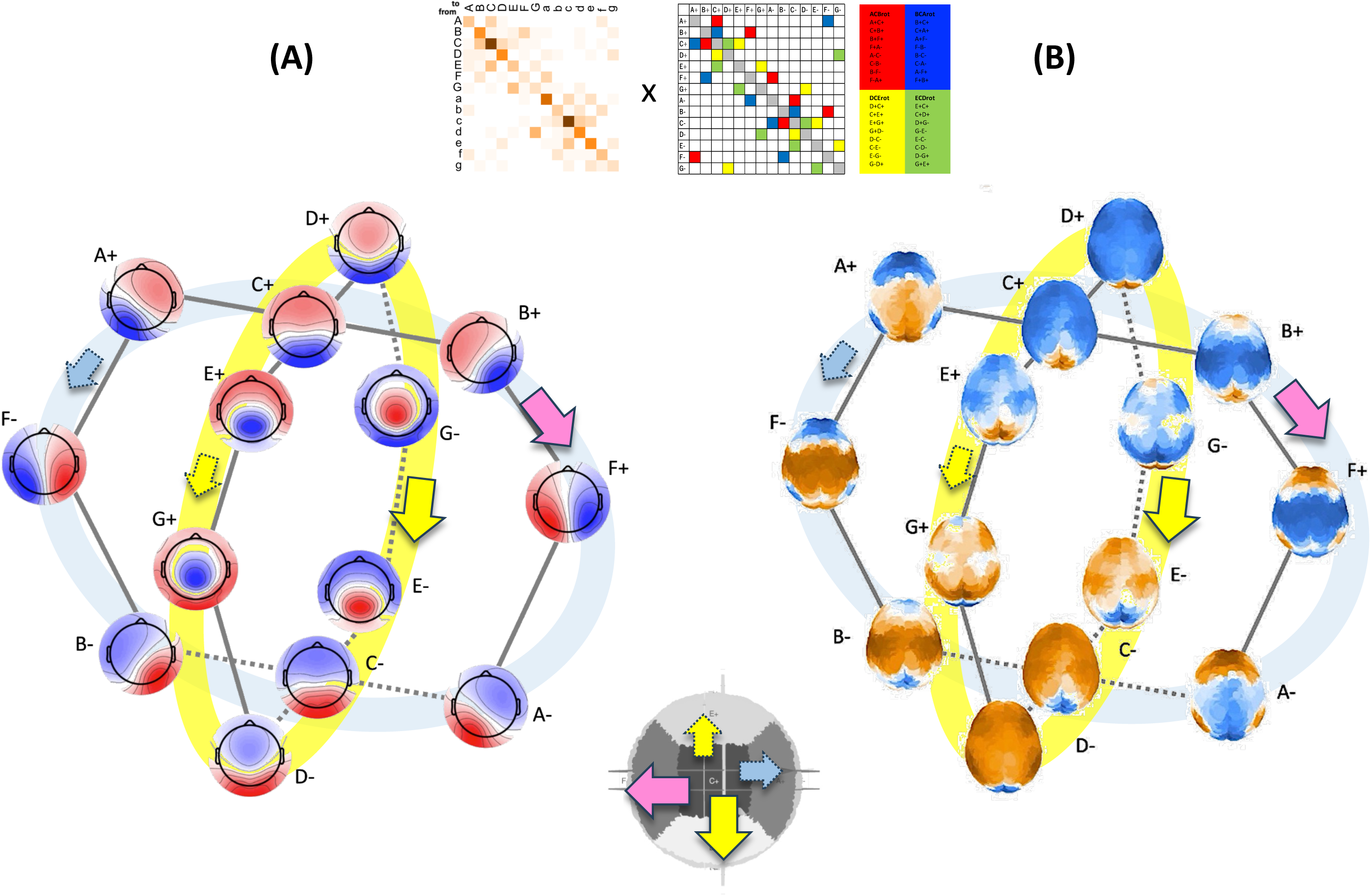
Topolo-geometrical state transitions (probability matrix) imply rotational trajectories. (A) For EEG state-dynamics and rotational trajectories. (B) For fMRI state-dynamics and rotational trajectories.

**Figure 7.**
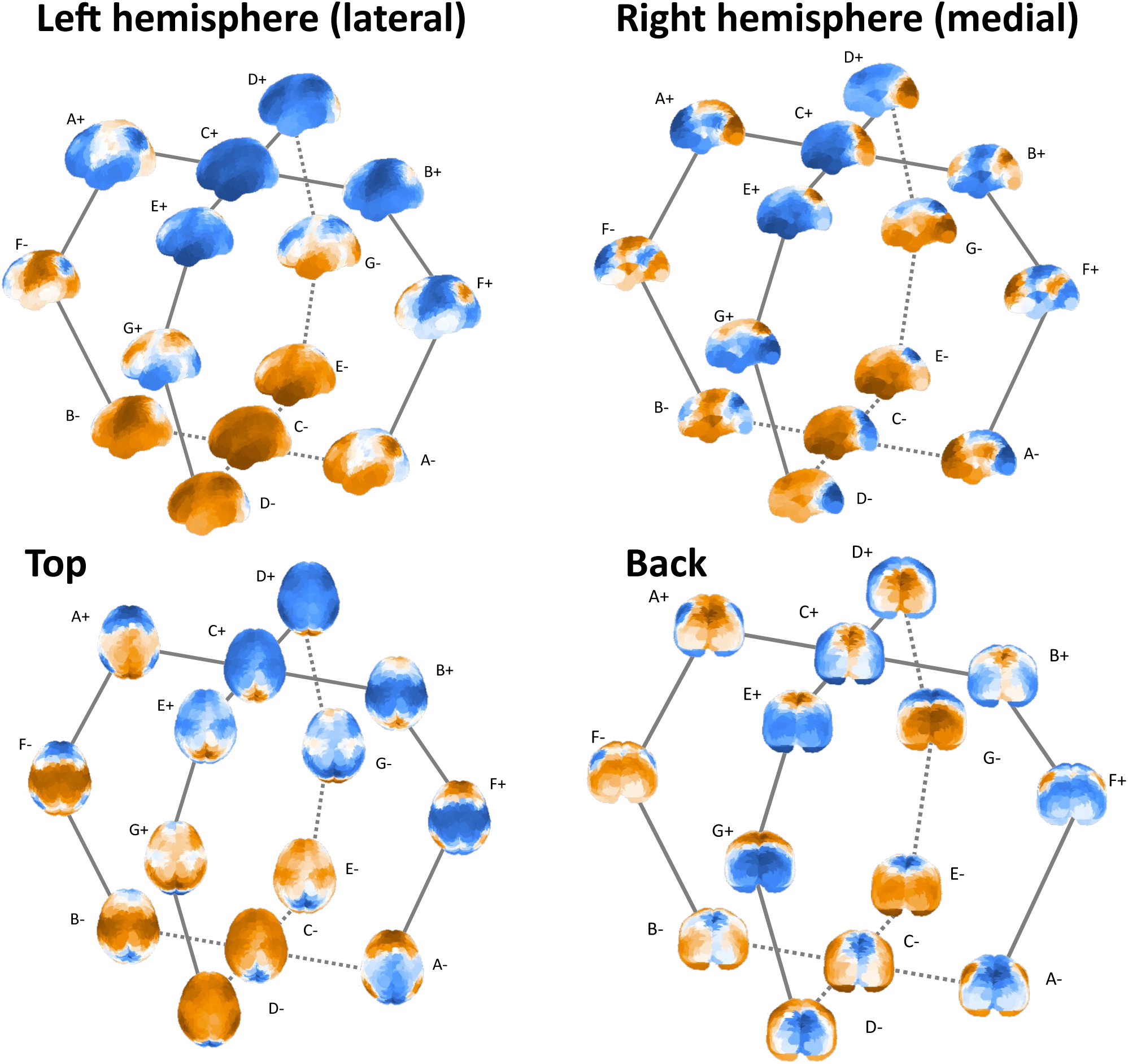
The ortho-diagonal spatial basis of the spherical state space (xyz = CFG) and their children (ABDE) in fMRI.

Figure 8 presents the central idea that an individual whole-brain neural state in EEG or fMRI can be represented as a point (x, y, z) in a common three-dimensional state space, and that this position directly determines its spatial pattern. As illustrated in the upper right, a “state” is therefore not treated as a discrete label, but as a location in a continuous space that is expressed as a scalp potential distribution in EEG or as a whole-brain activity pattern in fMRI. This idea is summarized by the equation shown in the center of the figure, in which a state pattern *v* is expressed as a linear combination of three basis patterns associated with the x-, y-, and z-axes. A point in the state space is thus not only an abstract coordinate, but also a set of coefficients that generates a concrete whole-brain activity pattern. In this sense, the three-dimensional embedding becomes a generative coordinate system rather than a purely descriptive one.

**Figure 8.**
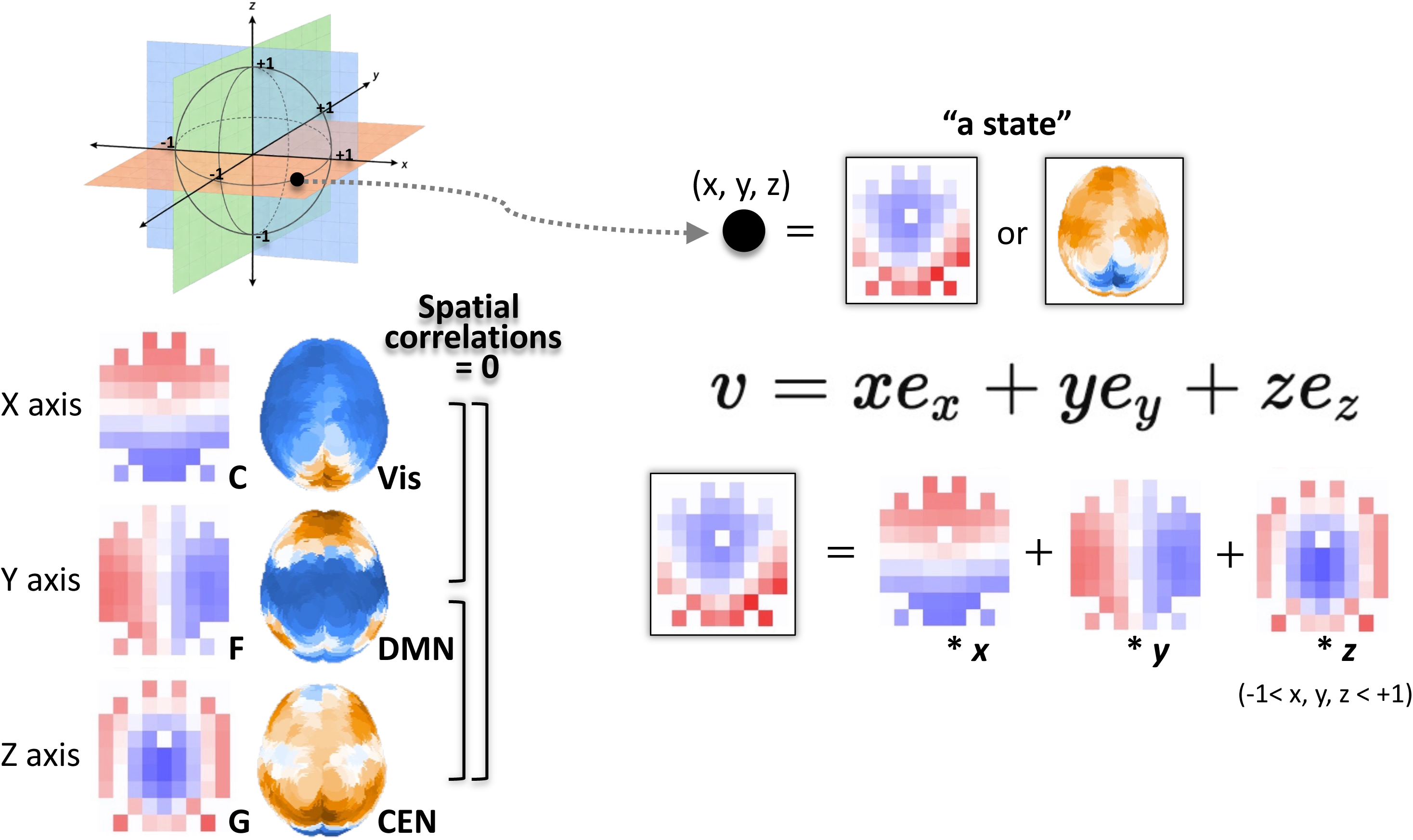
Ortho-diagonal state space produces the variation of the possible states as the results of the weighted linear sum of the three spatial basis.

The left side of the figure shows the basis patterns corresponding to each axis in both EEG and fMRI. In EEG, the three axes correspond approximately to C-like, F-like, and G-like scalp topographies. In fMRI, they appear as patterns resembling the visual (Vis), default mode (DMN), and central executive (CEN) systems. These three axes are treated as mutually orthogonal components with zero spatial correlation (see Fig. 3C and 5C), meaning that they form an independent basis for the state space. Any individual state can therefore be represented uniquely as a weighted sum of these basis patterns. The schematic in the lower right further clarifies this reconstruction principle. An EEG state pattern is expressed by weighting the x-, y-, and z-axis components by their respective coordinates, with each coefficient varying continuously within the range −1<x,y,z<+1 since the distance is based on the correlation. As the position of the state moves within the space, the balance among the three basis patterns changes accordingly. Individual states are therefore not “selected” from a discrete set, but generated as continuous combinations on a shared basis. A key implication of this figure is that EEG and fMRI states are not treated as separate modality-specific templates, but as different manifestations of the same underlying state vector defined by a common geometric principle. The same coordinate (x, y, z) is observed as a scalp topography in EEG and as a whole-brain network pattern in fMRI. This supports the view that the GeoMS framework is not merely an empirical clustering scheme, but a modality-independent coordinate representation built on shared basis axes.

More broadly, this approach replaces the conventional view of brain states as separate categories with a vector-based view (Wadia et al., 2026) in which each state is a position on a continuous manifold. Differences between states are therefore quantified not as category differences, but as differences in their coordinates along the basis axes. From this perspective, state transitions are understood as continuous movements through a common basis space rather than switches between templates. To illustrate this property, we developed several applications (see below for details). Figure 9 illustrates how the three-dimensional basis of the state space can be used not only to describe EEG/fMRI states, but also to decompose, reconstruct, and interpret them. In (A), states are treated as coordinates on a sphere at different spatial scales—channel, voxel, and ROI level—and the corresponding brain activity patterns are generated from those coordinates. In (B), Neurosynth annotation maps are embedded into the same spherical space, allowing the axes of the state space to be related to cognitive and functional concepts.

**Figure 9.**
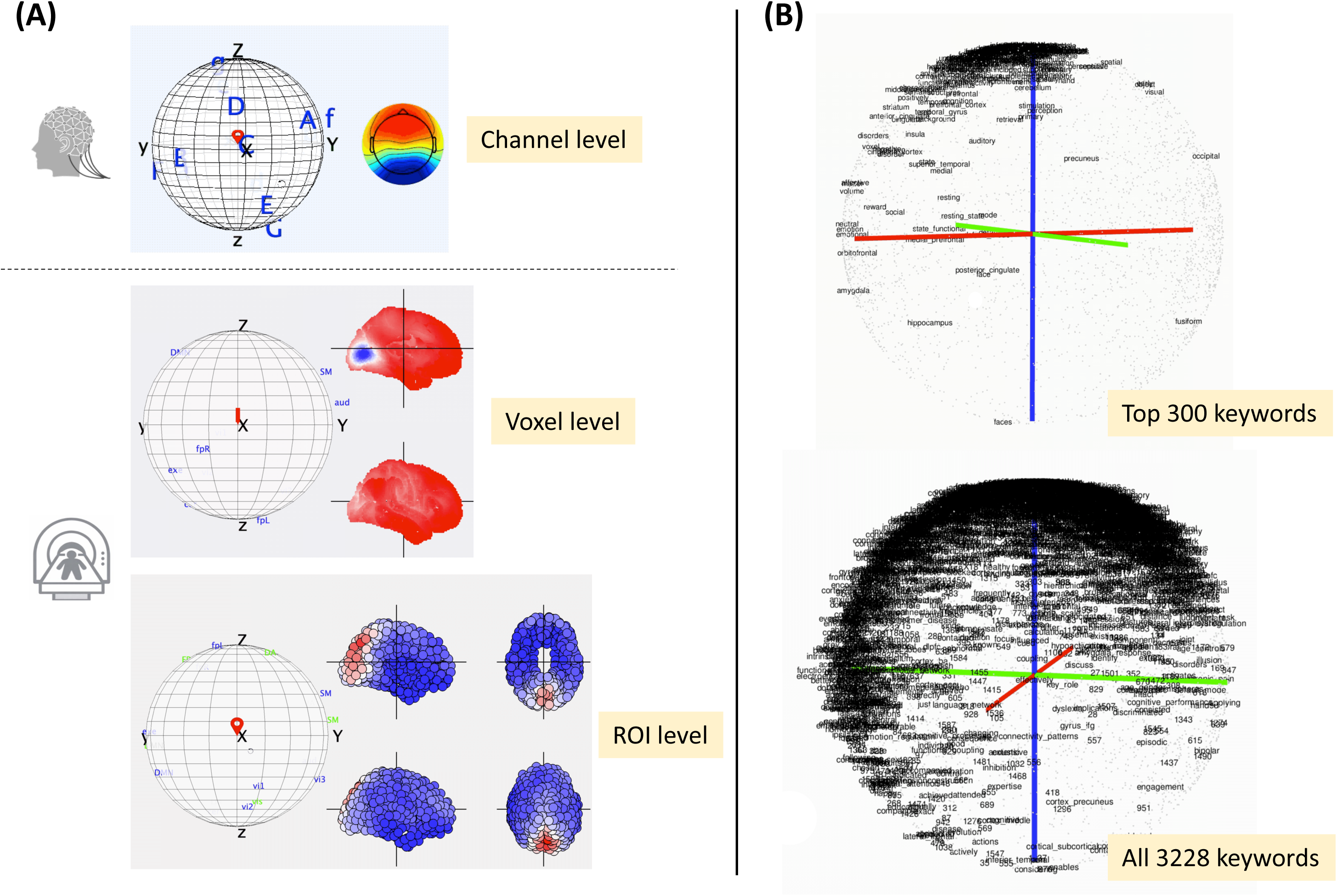
Application of the spatial basis for state decomposition and reconstruction. (A) The real-time generation of the EEG/fMRI states as an interactive app (terrestrial globe). (B) The spherical embedding of Neurosynth annotation maps.

In (A), the left column shows that the same spatial basis can be used across multiple representational scales. In each panel, a point on the sphere specifies a state coordinate, and that coordinate immediately generates the corresponding pattern on the right: an EEG scalp topography at the channel level, an fMRI whole-brain voxel pattern at the voxel level, or an ROI-level pattern at the coarser regional level. This makes explicit that states are not selected from a fixed library of templates, but are reconstructed continuously from their coordinates along the basis axes. At the channel level, the figure shows how EEG scalp maps vary smoothly as the position moves across the sphere, indicating that neighboring points in the state space correspond to neighboring topographic patterns. At the voxel level, the same geometric principle is applied to fMRI, where whole-brain spatial patterns are generated directly from the same type of coordinates. At the ROI level, the same reconstruction remains possible even after coarse-graining the data into region-based representations. Taken together, these panels show that the proposed basis is not tied to a single resolution or modality, but functions as a shared representational framework across scales. In that sense, the state space is not merely an analytic summary, but a generative coordinate system that can actively produce and manipulate brain-state patterns. The globe-like interface shown in the figure further suggests a practical implication of this framework (see the Supplementary Information for the demo application and accompanying video). Because moving to a new position on the sphere immediately changes the generated EEG or fMRI pattern, the state space can serve as both an abstract model and an intuitive interface for exploration, visualization, and interactive control. This gives the framework a concrete operational value beyond static representation.

In (B), the same spherical space is used to embed Neurosynth-derived annotation maps. The upper panel shows the distribution of the top 300 keywords, and the lower panel shows the embedding of all 3,228 keywords (currently). The colored axes represent the principal basis directions of the state space, and each keyword is positioned according to how strongly its annotation map aligns with those directions. This means that the same coordinates used to reconstruct brain states can also be used to locate functional terms within a common geometric framework. An important implication of this panel is that the axes of the state space are not merely abstract mathematical components, but are systematically related to known cognitive, perceptual, and behavioral concepts. Terms related to vision, audition, motor function, memory, emotion, and social cognition, for example, occupy different regions of the sphere, suggesting that the basis axes capture not only geometric variation in brain states but also aspects of their functional meaning. The top-300 map makes the overall organization easier to see, while the full 3,228-keyword embedding shows that this structure remains visible even in a much denser semantic space.

These analyses indicate that the spatial basis serves at least three roles. First, it provides a basis for decomposing EEG and fMRI states into common components. Second, it acts as a generative model, allowing state patterns to be reconstructed from arbitrary coordinates. Third, it supports semantic annotation, by embedding functional vocabulary into the same space and thereby linking geometry to interpretation. Thus, the proposed approach presents the state space as a unified framework rather than only the result of dimensionality reduction for generating, manipulating, and interpreting whole-brain neural states. Conventional template-based approaches may list representative patterns, but they do not easily allow continuous interpolation between them, nor do they naturally connect different modalities or functional vocabularies within a shared coordinate system. Our application shows that a spatial-basis framework can integrate all of these functions within a single geometric space. More strongly stated, the main message of this is that the three-dimensional basis is a generative and annotatable representational space, not just a visualization aid. Coordinates on the sphere serve as coefficients for reconstructing EEG/fMRI patterns at channel, voxel, and ROI levels, while the embedding of Neurosynth maps gives those same coordinates cognitive-functional meaning. The spatial basis therefore functions as a modality-independent, scale-independent, and semantically extensible common coordinate system, rather than only a low-dimensional compression of states.

Finally, we examined a practical application of this approach to biological individual differences in age, sex, and handedness. Figure 10 compares state-transition metrics between a younger group (blue, N=130) and an older group (yellow, N=55) in the LEMON dataset (the transition counts calculated at the original sampling rate of 2500 Hz), using GfpMS14 (EEG), GeoMS14 (EEG), and GeoMS14 (fMRI). The top row shows rotational transitions, the bottom row shows self-recursive transitions, and error bars indicate standard error (±SE). In the left column (GfpMS14), four rotational transition measures (ACBrot, BCArot, DCErot, ECDrot) and state-wise self-recursive counts are shown separately for the eyes-closed and eyes-open EEG resting conditions. Although some age-related differences are visible, they are generally modest and not especially consistent across metrics or conditions. Thus, the conventional template-based definition detects some aging effects (Kashihara et al., 2025), but these effects appear relatively weak and dispersed. In the middle column (GeoMS14, EEG), the same EEG data are analyzed using the topolo-geometric landmark approach. Here, age-related differences are more pronounced and more consistent than in GfpMS14. In particular, older participants tend to show higher values for several rotational transition measures across both eyes-closed and eyes-open conditions. Differences in self-recursive transitions are also clearer, suggesting that aging is associated with altered circulation among states and a redistribution of which states are more likely to persist. Relative to the template-based solution, GeoMS14 therefore appears to capture aging-related changes in EEG dynamics more sensitively.

**Figure 10.**
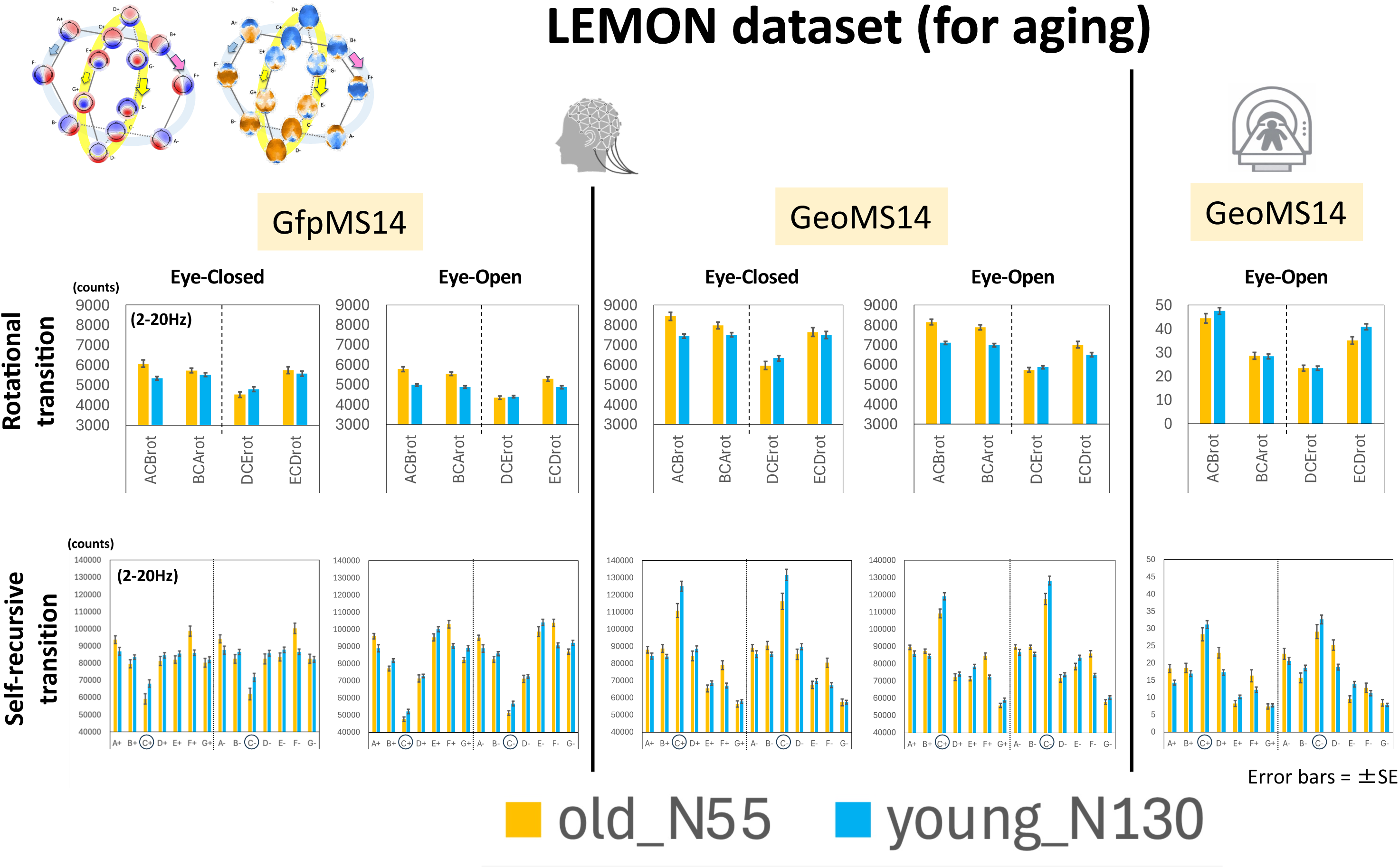
Rotational and self-recursive transitions for distinguishing between the old and the young groups. Note: Identified rotational (upper) and self-recursive (lower) differences in aging depend on the templates and the modality.

The right column shows the fMRI version of GeoMS14, computed from the eyes-open fMRI resting data (LEMON). As in EEG, age-related differences are observed in both rotational and self-recursive measures, although the absolute scales are much smaller because of the lower temporal resolution and shorter state sequences in fMRI. Even so, specific rotational modes and self-recursive states still differ between younger and older participants, indicating that the sensitivity of GeoMS14 to aging-related dynamics extends beyond EEG to fMRI. Two points are especially important. First, for the same aging contrast, GeoMS14 in EEG appears to extract group differences more systematically than GfpMS14. Second, a similar tendency is also visible in GeoMS14 for fMRI, suggesting that these transition-based measures may capture aging effects across modalities. This result indicates that aging is expressed as a reorganization of brain-state dynamics—that is, differences in how states rotate through the state space and how long particular states tend to persist. It further suggests that these dynamic changes are more clearly captured by GeoMS14, which preserves the geometry and directionality of the state space, than by the conventional template-based approach.

In addition, Figure 11 shows how rotational transitions based on GeoMS14 reflect differences related to sex and handedness in the EEG608 dataset. The upper panels compare the frequencies of four rotational transitions (ACBrot, BCArot, DCErot, ECDrot) across eyes-closed and eyes-open conditions, while the lower panels compare directional bias measures derived from their differences across frequency bands. In the upper row, rotational transition counts are compared across four groups: female left-handed, male left-handed, female right-handed, and male right-handed. Some group differences are visible in the individual rotation counts, but their interpretation is somewhat diffuse when each measure is considered separately. For example, differences between ACBrot and BCArot, or between DCErot and ECDrot, are more naturally understood not simply as increases or decreases in total rotation, but as asymmetries between opposite directions of rotation. To capture this more directly, we define two directional bias measures. The first is the horizontal directional bias, defined as ACBrot − BCArot, and the second is the vertical directional bias, defined as ECDrot − DCErot. These measures quantify not just how often rotations occur, but which direction is preferred within the state space. In the lower bar plots, these two directional biases are compared across delta (1–4 Hz), theta (4–8 Hz), alpha (8–12 Hz), beta (15–30 Hz), 2–20 Hz, and 1–30 Hz bands. The biases are especially prominent in the alpha and beta ranges, suggesting that directional effects are more pronounced in these functionally important mid-frequency bands than in slower rhythms. Across groups, the vertical bias appears to separate the groups more strongly than the horizontal bias, and the alpha range is visually highlighted as particularly informative. The bottom line plots further show these directional biases as continuous functions across 1–30 Hz, revealing that group differences depend not only on broad frequency bands but also on finer spectral positions. Some groups and conditions show stronger vertical bias around 8–10 Hz, whereas others show stronger horizontal bias at higher frequencies, indicating that each has a distinct frequency-dependent profile of rotational anisotropy.

**Figure 11.**
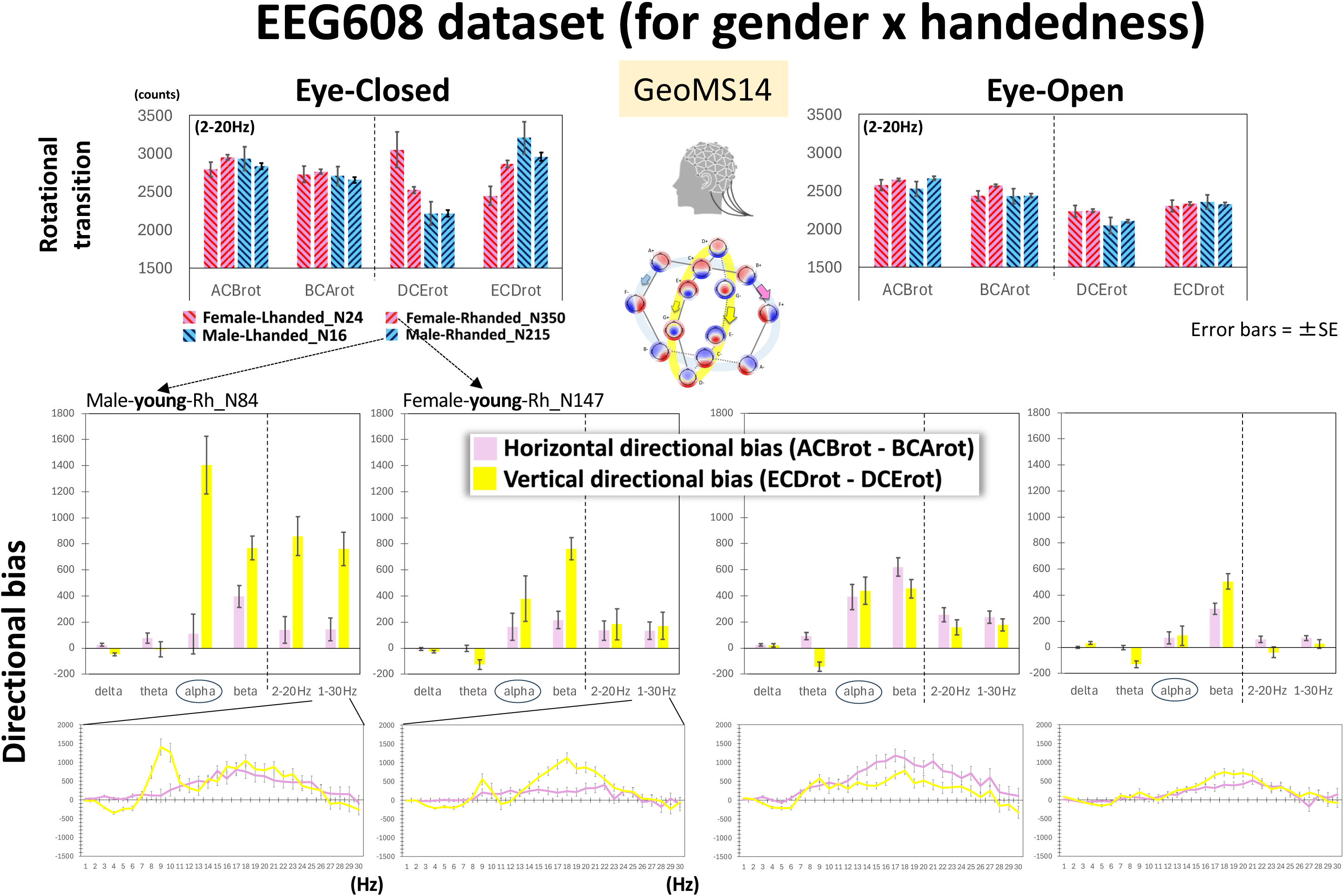
Rotational direction bias characterizes the gender and/or handedness groups. Note: Rotational transition (upper) reveals the direction bias between ACB-BCA and ECD-DCE. However, this balance differs among gender x handedness groups. The horizontal and vertical direction bias (lower) emphasizes the gender difference.

Differences related to sex and handedness emerged more clearly as rotational-direction biases than as simple differences in state frequency or total transition count (e.g., Kashihara et al., 2025). If the four groups differed only in overall activity level or total amount of rotation, subtracting opposite rotations would reduce group differences. Instead, the differences become clearer after this subtraction, suggesting that the groups are distinguished less by how much they transition than by how their state trajectories are directionally organized. Overall, the result suggests that the effects of sex and handedness may be expressed as a sex × handedness interaction pattern rather than as independent main effects, with each subgroup showing a distinct directional bias profile. In this sense, GeoMS14-based rotational measures provide a sensitive way to characterize biological individual differences in EEG dynamics.

## Discussion

The present study redefined the template problem in EEG microstate analysis as a problem of the topographic structure rather than simply one of selecting representative maps through clustering of the state space spanned by scalp potential maps. On this basis, we proposed a framework in which templates are not derived from GFP-peak, polarity-invariant clustering, but are instead rediscovered as discrete representatives of dominant axes embedded in a broader state space constructed from mutual similarities among whole-brain states. The results showed that these redefined templates, or “landmarks”, achieved better performance than conventional templates. This finding is not merely a technical improvement of one algorithm over another. It suggests that the very definition of states in microstate analysis and other similar approaches may need to be reconsidered.

Conventional microstate analysis has been highly successful as a means of coarse-graining continuous brain dynamics into a small number of discrete states. At the same time, however, it has long carried a structural limitation: nearly all downstream measures, including duration, occurrence, coverage, and transition probability, depend strongly on the initially estimated templates. The present findings suggest that this limitation may arise not only from implementation noise but also from defining templates themselves as cluster centroids. In the conventional framework, templates are obtained as local outcomes of clustering and are therefore susceptible to variation arising from initialization, sampling bias, preprocessing choices, and the selected number of clusters. By contrast, the landmarks introduced here are derived from the global geometric structure of the state space and represent dominant directions embedded in the overall topography. Their improved performance suggests that such globally constrained nodes may provide a more stable and valid basis for state definition than locally defined cluster centers.

From this perspective, one of the central contributions of the present work is the conceptual shift from “templates” to “landmarks.” Conventional templates are defined as averages or centroids within clusters and therefore represent the outcome of grouping similar observations together. Landmarks, in contrast, are defined as topographic nodes associated with dominant axes of state space and therefore reflect directions that organize the space as a whole. This distinction is fundamental for any attempt to discretize a continuous process. Discrete states should be defined by positions that capture the structural backbone of the underlying dynamics. Our findings support the possibility that, in microstate analysis, the most useful nodes are the landmarks that best span the geometry of the state space, with C± appearing as particularly central, global attractor-like landmarks (Croce et al., 2020).

The present study also prompts a reconsideration of polarity. In conventional polarity-invariant clustering, voltage maps with opposite signs are treated as instances of the same template, thereby simplifying the analysis. However, this assumption may also remove, in advance, an important geometric property of scalp maps when they are interpreted as directional vectors in a continuous state space: namely, their antipodal relationship. In the present approach, polarity is retained as a natural geometric relation within state space. The finding that such polarity-preserving representations yielded better performance suggests that, at least for some forms of brain-state description, ignoring polarity a priori may have led to a loss of information. This does not imply that polarity-sensitive analysis will be preferable in every context, and the physiological interpretation of sign inversion remains to be examined with caution. Nevertheless, our results indicate that the conventional practice of removing polarity should not be treated as the only justified option (Kashihara et al., 2025; Tsubaki et al., 2026).

Another implication concerns the longstanding issue of choosing the number of clusters, k (Michel & Bréchet, 2026). In conventional microstate analysis, there has been persistent tension between the historical assumption of four or five canonical microstates and the practical need to optimize k in a data-driven manner. In the present framework, landmarks are determined by extracting dominant axes from the geometry of the state space. As a result, the influence of the k problem can, at least in principle, be relativized. The number of landmarks is no longer framed primarily as the number of clusters to be found, but as the number of principal directions required to describe the structure of the continuous space. This shift expands microstate analysis from a clustering-centered methodology into one grounded in state-space geometry.

Our “practical” findings suggest that the effects of eyes-open/eyes-closed, sex, handedness, and aging in EEG are not merely expressed as differences in static spectral power (Han et al., 2025; Martin et al., 2025), but can also be characterized as directional biases in rotational state transitions within the neural state space (Kashihara et al., 2025). Previous EEG studies have shown that sex differences are distributed across a wide range of measures, including absolute and relative spectral power, hemispheric asymmetry, and functional connectivity, rather than being confined to a single frequency band or scalp region. In children, boys and girls already exhibit widespread differences across these EEG features (Modarres et al., 2023), whereas in adults, sex differences become more pronounced with aging, particularly in the low-alpha and low-beta ranges (Han et al., 2025). Our results are broadly consistent with these earlier observations, but extend them by suggesting that such group differences may also be understood at the level of geometric trajectory, namely, as asymmetries in the preferred direction of state transitions.

In the EEG608 dataset, the four sex-by-handedness groups showed only partially separable differences when the individual rotational transitions themselves were examined. In contrast, these differences became much clearer when transitions were collapsed into directional indices, namely the horizontal directional bias (ACBrot − BCArot) and the vertical directional bias (ECDrot − DCErot). This pattern suggests that the essential group difference is the direction in which the trajectory tends to circulate within the state space. In this sense, the sex effect appears not only as a difference in signal magnitude, but as an asymmetry in dynamical flow. This pattern was further modulated by handedness, implying that handedness should not be treated merely as a nuisance covariate. Rather, it may act as a factor that reshapes the manifestation of sex-related asymmetry in large-scale EEG dynamics (Davidson et al., 1976; Galin et al., 1982; Glass et al., 1984; Ocklenburg et al., 2019).

The LEMON dataset further supports this interpretation from the perspective of aging. Differences between the young and old groups were observed not only in rotational transitions but also in self-recursive transitions (Kashihara et al., 2025). This combination is important, because it indicates that aging may alter not only the dwell structure of metastable states, but also the preferred circulation of trajectories across states. In other words, the aging effect observed here cannot be reduced to a simple increase or decrease in transition counts; instead, it points to a trajectory-level reorganization of neural dynamics. This interpretation provides a potentially useful bridge to previous reports that aging amplifies sex differences in low-alpha and low-beta oscillations (Han et al., 2025). Rather than viewing those findings only as local spectral effects, our results raise the possibility that such band-limited differences reflect a more global change in the directional organization of state-space dynamics. Taken together, these findings support the view that sex, handedness, and aging are reflected in the non-equilibrium geometry of neural state transitions as well as in static EEG and fMRI descriptors of neural state transitions. The present results therefore argue for a shift in emphasis from asking which state occurs more often toward asking how neural activity flows through the state space and whether that flow is directionally biased across groups. Such a perspective may help integrate conventional EEG findings with state-transition-based approaches, and may provide a more unified view for understanding group differences in large-scale neural dynamics.

Several limitations should be acknowledged. First, although the present study demonstrated improved performance of landmarks over conventional templates, the generality of this advantage across EEG conditions, participant populations, and electrode configurations remains to be established. In particular, whether landmarks remain stable across recording setups and preprocessing pipelines will be critical for the broader claim that they may support more consistent comparisons across studies and across EEG caps. Second, although landmarks reflect axes of state space, they should not be interpreted as directly corresponding to unique physiological generators or functional units. Their neurophysiological meaning therefore requires further investigation, ideally through source analysis or multimodal recordings. Third, while the present framework redefines how templates are constructed, its relationship to key themes in the microstate literature—such as transition dynamics, inter-individual variability, and disease specificity—still needs to be examined in future work.

Despite these limitations, the implications of the present study are clear. State transition-based analysis has traditionally developed as a framework in which time-series measures are compared after templates have been taken as given. The present results show that this point of departure itself can be reformulated in a more principled way. Templates need not be understood as cluster centroids extracted from a particular dataset, but may instead be interpreted as landmarks rediscovered from the topology of the state space. By relocating the concept of the template from a local clustering result to a geometrically grounded axis of neural-state organization, the present framework preserves the original strength of microstate analysis—its ability to coarse-grain rapid continuous dynamics—while offering a more stable and potentially more unified definition of nodes. In this sense, the present study does extend conventional microstate analysis its conceptual foundation in a topological and geometric direction, thereby opening a path toward a next-generation framework that may better support comparisons across studies and across recording systems.

## Conclusion

The introduction of a topolo-geometric or landmark-based state definition was critical for uncovering the present findings. Unlike conventional template-based approaches, which mainly emphasize state occurrence or duration, the landmark framework preserved the global geometry of the state space and thereby exposed systematic directional asymmetries in neural trajectories. As a result, sex, handedness, and aging effects became interpretable as differences in static EEG features and in the preferred flow structure of whole-brain state dynamics. This suggests that the key contribution of the landmark approach is not simply methodological refinement, but the ability to reveal an otherwise hidden dynamical organization of group differences in EEG.

## Supplementary Information

The demo application and accompanying video are available online: https://www.dropbox.com/scl/fi/wsnwt0jo77s08tdz800sb/Asai2026_SI.zip?rlkey=4v0u9whmeip65rm2sn8×6s67t&st=ikio4f39&dl=0. The application is provided as a MATLAB P-file; the source code itself is not yet publicly available. On a Mac, it should be possible to run it by right-clicking the file and selecting “Open With → MATLAB.” However, we do not guarantee operation across different MATLAB versions or environments, nor do we provide technical support. Please also note that this is a work-in-progress version and may differ from the latest version. We have summarized derivative versions and examples of use in the accompanying video, which we encourage readers to view as well.

## Acknowledgements

During the preparation of this manuscript, the authors received assistance from ChatGPT 5.4 and 5.5 for literature review, programming, drafting of the text, and English-language revision. The authors critically reviewed and edited all outputs and are solely responsible for the accuracy, interpretation, and final content of the manuscript.

## Declaration of Competing Interest

The authors declare no competing interests.

## Author Contributions

TA conceived and designed the study, collected the data, performed the analyses, interpreted the results, and drafted the manuscript. SK and SC contributed to all aspects of the study, with SK providing particular expertise and support for the EEG analyses and SC for the fMRI analyses. All authors read and approved the final manuscript.

## Funding

This work was supported by the JST Moonshot R&D Program (Grant Number JPMJMS2291) to TA, SK, and SC; AMED (Grant Number JP25wm0625219) to TA and SC; and AMED (aGrant Number JP25wm0625402) to TA.

## Notes

https://www.dropbox.com/scl/fi/wsnwt0jo77s08tdz800sb/Asai2026_SI.zip?rlkey=4v0u9whmeip65rm2sn8x6s67t&st=ikio4f39&dl=0

### Competing Interest Statement

The authors have declared no competing interest.

